# The accumulation of colibactin intermediates does not affect the growth and morphology of *pks*^+^ *Escherichia coli*

**DOI:** 10.1101/2020.04.17.047712

**Authors:** Yermary Morales-Lozada, Ramón Gómez-Moreno, Gabriela Báez-Bravo, Iraida E. Robledo, Dámaris Suazo-Dávila, Carlos R. Cabrera, Abel Baerga-Ortiz

## Abstract

Colibactin is a natural product made by numerous strains of *E. coli* that harbor the *pks* genomic island. The deletion of one of the genes within the *pks island,* the peptidase clbP, has been found to disrupt the maturation of colibactin, thus promoting the accumulation in the periplasmic space of numerous biosynthesis intermediates, some of which have been characterized chemically. To date, no one has reported the effect of such an accumulation of intermediates on the cellular morphology of the producing *E. coli* bacterium. In this report, we describe the scanning electron microscopy (SEM) images of numerous clinical isolates of *E. coli* harboring the *pks island*, collected from Puerto Rico hospitals. We have observed that the wild type isolates that harbor the *pks island* display lesions on the bacterial envelope surface. These lesions are absent in isolates lacking the *pks island*. To determine whether this phenotype is associated with colibactin production, we deleted the *clbP* gene from the extraintestinal pathogenic *E. coli* strain IHE3034, thus disrupting its ability to make colibactin. The wild-type IHE3034 displayed a spherical shape with no envelope lesions, and was practically indistinguishable from the Δ*clbP* deletion mutant. To our knowledge, this work provides the first SEM images of a pks deletion mutant.

**Importance:** The *pks* genomic island has been linked to the promotion of DNA damage and colorectal cancer, through the production of genotoxic compound colibactin in some strains of *E. coli.* While much is known about the mechanism of colibactin toxicity once it enters the mammalian cell, the prior steps leading to colibactin secretion or translocation from the bacterial cell, remain unclear. Here, we report high-resolution electron microscopy images of *E. coli* IHE3034 strain, a known colibactin producer, and a deletion mutant that is known to accumulate colibactin intermediates. The images reveal a predominantly spherical morphology that is unaffected by the accumulation of colibactin precursors and intermediates.

## Introduction

Colibactin is a genotoxic natural product made by some strains of Escherichia coli and other gram-negative bacteria, that carry the *pks island* gene cluster (1, 2). Among the genes encoded in the pks island are numerous polyketide synthase / non-ribosomal peptide synthases (PKS/NRPS) as well as other auxiliary proteins that work in concert for the production, processing and trafficking of the elusive colibactin (1, 3) which has been found to cause effects on the infected mammalian cell, including DNA damage, cell cycle arrest and megalocytosis (1, 4). Such genotoxic effects have been associated with an increase in the risk of colorectal cancer, especially in patients with inflammatory bowel disease (IBD)-associated cancer (4–6).

The biochemical routes for the synthesis of colibactin have been almost completely delineated and a mechanism of genotoxicity has been identified. It has been shown that several PKS/NRPS enzymes synthesize the inactive precursor, pre-colibactin, inside the bacterial cell.(7–9) The pre-colibactin is subsequently translocated to the periplasmic space by a transporter (ClbM)(9), activated by a peptidase (ClbP) (7, 8, 10, 11), and finally exported out of the bacterial cell (Figure 1). Exactly how the product reaches the nucleus of the affected mammalian cell, is still not known. However, once in the nucleus, colibactin can form DNA cross-links and double-strand breaks that activate much of the DNA damage responses while also inducing cell cycle arrest (12–14). Recently, the groups of Nougayréde and Crawford, independently showed how specific covalent adducts between DNA and colibactin can be formed (13, 15). Furthermore, at around the same time, Balskus group found evidence of adduct formation in a germ-free mice infected with *pks^+^ E. coli,* showing that the consensus mechanism of colibactin toxicity also operates *in vivo* (14).

**Figure 1.**
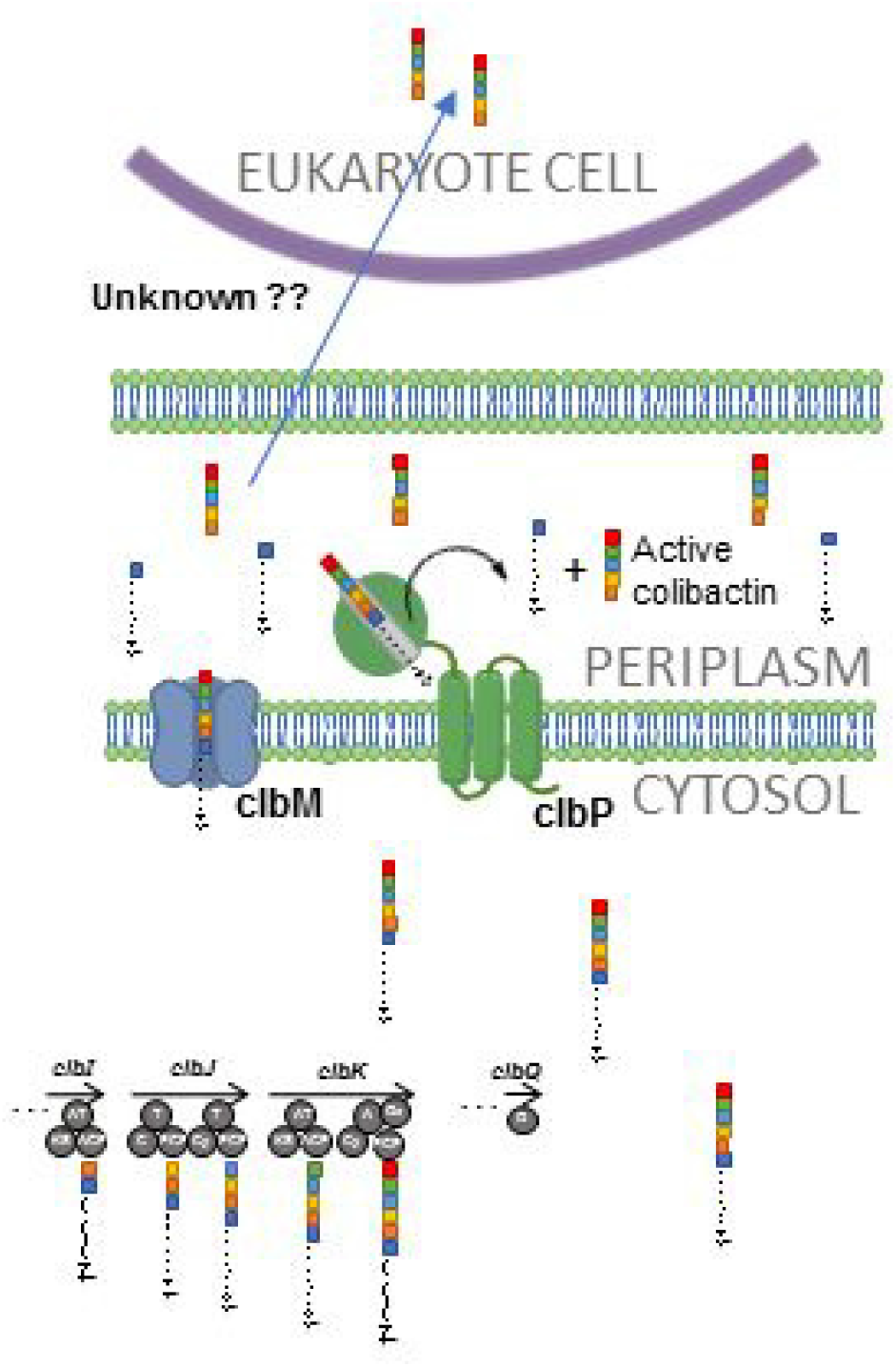
Schematic representation for the production and genotoxic effect route for colibactin. PKS/NRPS enzymes synthesize pre-colibactin inside the bacterial cell. The pre-colibactin is subsequently translocated to the periplasmic space by transporter (ClbM), activated by a peptidase (ClbP), and finally exported out of the bacterial cell toward the target cell by an unknown mechanism.

Colibactin and its intermediates are notoriously difficult to isolate and detect under standard conditions for chemical extraction and analysis (1). In fact, the production of adequate amounts of pre-colibactins, and other intermediates for chemical elucidation, has required the deletion of genes along the biosynthesis and processing route, so as to cause an accumulation of intermediates that would render them amenable for extraction and processing. For example, the deletion of the gene clbP was a key development in identifying the first colibactin intermediates through their accumulation in the periplasmic space (10, 16). This accumulation and processing of compounds in the bacterial periplasmic space would be expected to cause damage or morphological changes to the bacterial envelope, resulting in a microscopic footprint of production.

This work is an attempt to document the effect of the accumulation of colibactin intermediates on the morphology of colibactin-producing *E. coli* cells by scanning electron microscopy (SEM). Although we initially detected membrane lesions on the surface of clinical isolates *of E. coli* harboring the *pks* genomic island, the deliberate genetic disruption of the island did not result in measurable morphological changes against wild type.

## Results

### SEM images of pks^+^ and pks^−^ clinical isolates

We were initially driven to study the effect of colibactin production and accumulation on *E. coli* morphology, from examining the scanning electron micrographs (SEM) presented here for several clinical isolates of *pks^+^* and *pks^−^* bacteria from a local strain repository at the University of Puerto Rico Medical School (see Figure 2). From these electron microscopy images, it was easy to identify lesions on the surface of the *E. coli* bacteria which were detected almost exclusively in the *pks^+^* isolates but absent in the *pks-* isolates. The apparent damage observed on the *E. coli* envelope of *pks^+^* strains was not different from the damage caused by the application of an antimicrobial peptide or by membrane-disrupting quantum dots (17).

**Figure 2.**
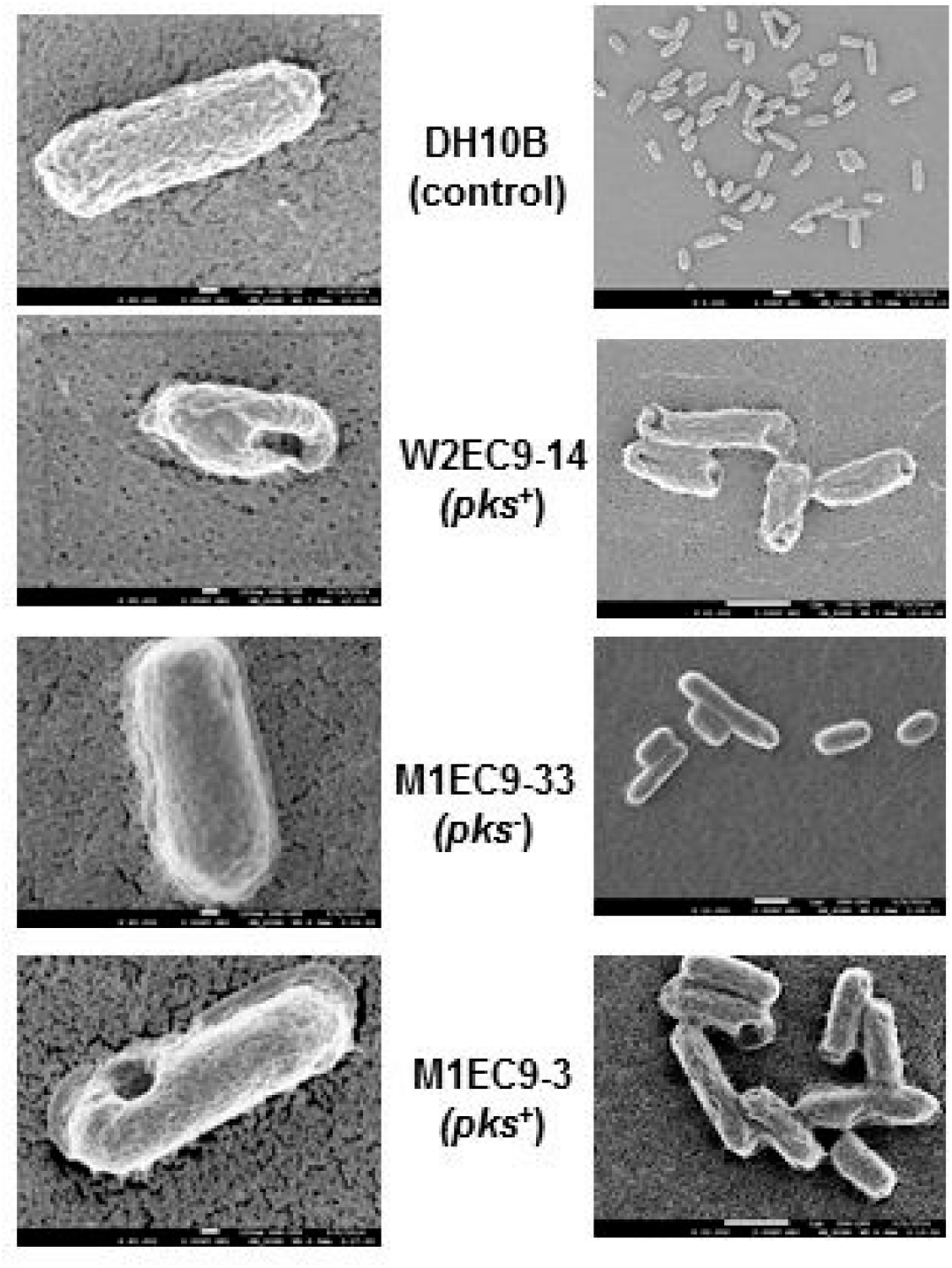
SEM images of *pks^+^* and *pks^−^* clinical isolates obtained from a local pathogen surveillance repository. The cells were grown in LB medium at 37 °C, overnight. Fixation, dehydration and gold coating of the samples were carried out on a glass surface coverslip.

### Deletion of the ClbP gene

In an effort to more specifically connect the observed membrane disruption shown in Figure 2 to the accumulation of colibactin intermediates, we deleted the *clbP* gene in the pathogenic strain IHE3034 by homologous recombination. The *ClbP* gene encodes a peptidase that is required for the hydrolysis of pre-colibactin, and whose absence abolishes the megalocytic phenotype in the infected mammalian cell and causes the accumulation of pre-colibactin and other intermediates (16). The deletion of *clbP* was confirmed by polymerase chain reaction (PCR) (see Figure 3a).

**Figure 3.**
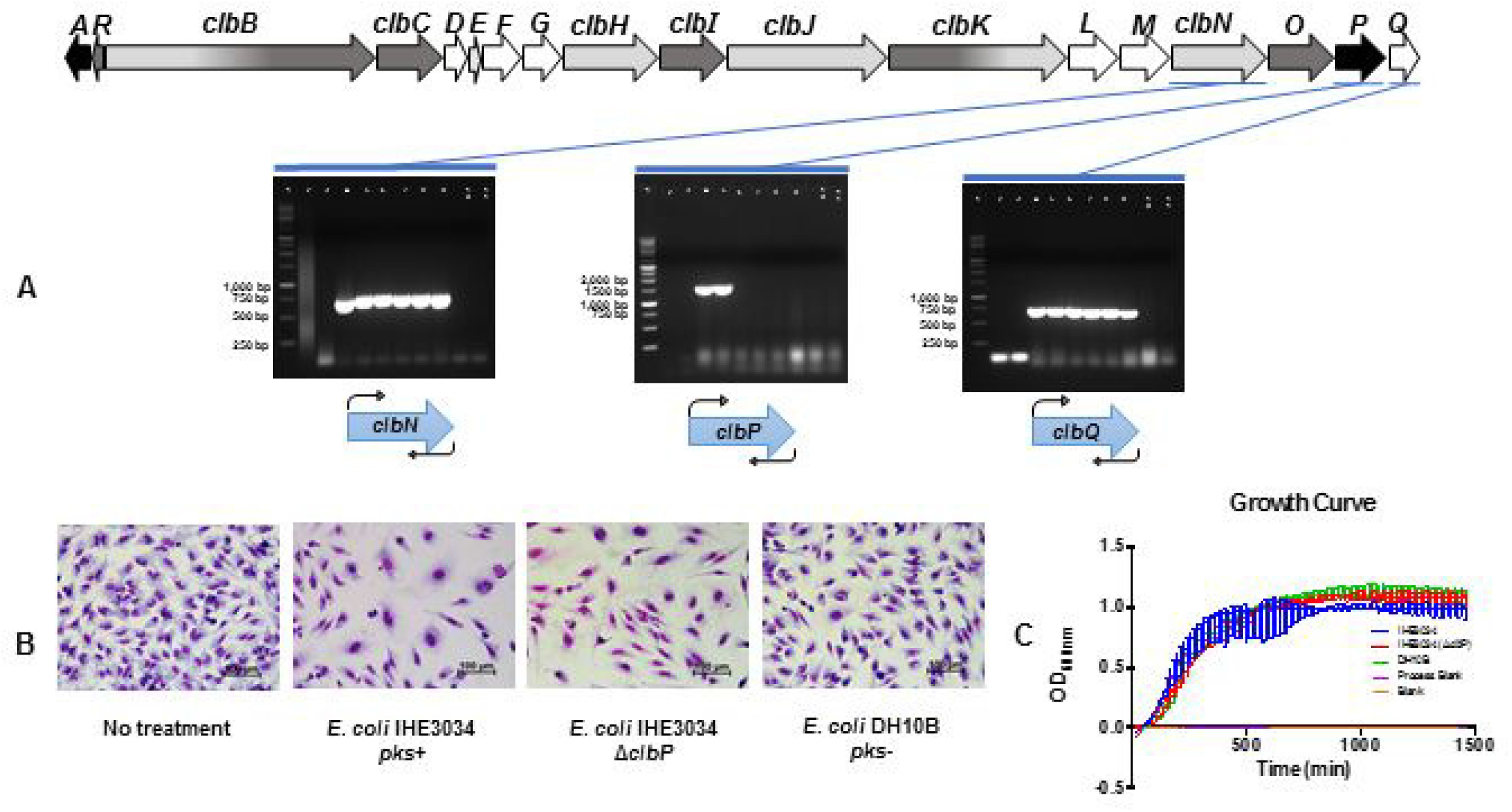
Deletion of the *ClbP* gene in *pks^+^ E. coli* strain IHE3034. **(A)** PCR confirmation of *clbP* deletion. (1) 1kb DNA standars, (2 and 3) PCR Blank (4 and 5) *E. coli* IHE3034 pks+ (6 to 9) *E. coli* IHE3034 Δ*clbP* (10 and 11) *E. coli* DH10B *pks*^−^. **B)** Giemsa staining of HeLa cell line after 72 hours of post-infection with IHE3034 *(pks^+^),* IHE3034 (Δ*clbP*) and DH10B (negative control) *E. coli* strains at 37 °C and 5 % CO_2_. Bacterial infection was for 4 hours at 37 °C and 5 % CO_2_. All optical microscope images were taken at 40X. **C)** Growth Curve of IHE3034 *(pks^+^),* IHE3034 (Δ*clbP*) and DH10B *E. coli* strains at 37 °C under constant shaking.

We also performed a cellular assay in which we infected HeLa cells with either wild type or Δ*clbP* IHE3034, to confirm the abolishment of the megalocytic phenotype associated with colibactin production. As expected, minimal or no megalocytosis was observed in the IHE3034 Δ*clbP E. coli* and in the DH10B *E. coli* (negative control strain), however, cells treated with *E. coli* IHE3034 *pks^+^* (wild type strain) showed the expected megalocytosis phenotype (Figure 3b). The deletion of *clbP* did not impair, nor did it enhance, the growth rate of IHE3034 *E. coli* (see Figure 3c).

### SEM images of wild type IHE3034 and IHE3O34ΔclbP E. coli

SEM images were collected for the IHE3034 wild type and mutant strains, following the exact same protocols employed for viewing the locally source strains. We expected that the deletion of the *clbP* gene would induce outer-membrane damage due to the accumulation of colibactin intermediates. To our surprise, the electron microscopy images of IHE3034 and IHE3034 Δ*clbP* revealed no morphological effect expected from the disruption of colibactin production (see Figure 4).

**Figure 4.**
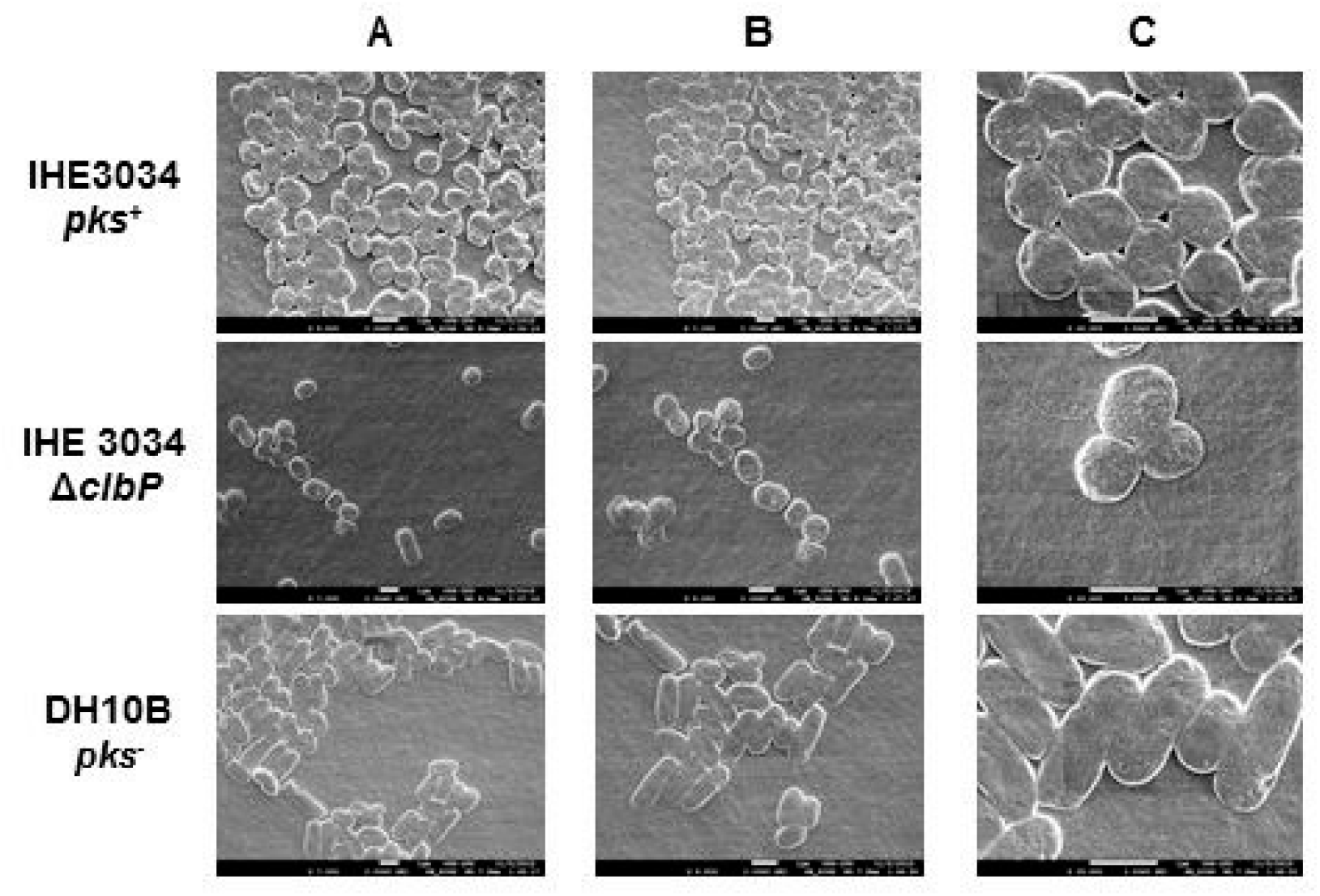
SEM images of IHE3034 *(pks^+^)* and its isogenic mutant strain Δ*clbP* vs DH10B (negative control), after 24 hours of incubation at 37 °C without antibiotic. Columns A, B and C are photos taken at 7,000, 9,500 and 25,000x of magnification, respectively.

## Discussion

Since its discovery reported in 2006, the *pks* genomic island and its resulting product colibactin have generated great interest (1). The postulated role of colibactin in colorectal carcinogenesis through DNA damage, provides a direct mechanistic connection between specific genes from the gut microbiota and cancer (4, 5, 18, 19). Not surprisingly, there have been numerous attempts to elucidate its biosynthesis (7, 8, 10, 11, 20–25), its chemical structure(10, 12, 16, 26–28), and its mechanism of genotoxicity (12–14). In many of these reports, there is a common observation of accumulation of colibactin intermediates in *E. coli* cells, enough to characterize chemically. These intermediates have been the basis of efforts to elucidate the structure of active colibactin (15).

Our work attempts to document the effect of such accumulation of the polyketide/peptide intermediates of colibactin within the periplasmic space of the producing bacteria. We expected that an accumulation of compounds, resulting from the deletion of the clbP peptidase, would cause a weakening of the bacterial envelope, which would lead to a deformation that could be readily detected by SEM. Although differences in the envelope integrity in some *pks^+^* wild-type clinical isolates were observed, these differences cannot be attributed to the presence or absence of the *pks* genes, since neither the wild type IHE3034 strain nor its mutant strain showed the expected morphological changes.

The fact that we were unable to see any differences in bacterial morphology or in envelope integrity resulting from the deletion of *clbP*, could suggest that the concentration of colibactin and its intermediates within the producing bacteria is actually quite low. This is not surprising since the production of enough pre-colibactin for NMR analysis has required the cultivation in scales of 60-200 liters of *E. coli* culture (27, 28). Moreover, it was recently proven that not all colibactin-maturing peptidases demonstrate the same bioactivity, which means that not all pks+ strains yield the same amount of its bioactive compound (29).

The IHE3034 strain of *E. coli* is a well-studied extraintestinal pathogenic (ExPEC) strain originally isolated from a newborn that developed meningitis (30). The fact that this strain is a clinical isolate, and hence contains various virulence factors(31), may contribute to our observation that in the infection assay of HeLa cells with IHE3034Δ*clbP*, the megalocytosis phenotype was not completely abolished as other groups have recently reported (16).

There are other reports describing how IHE3034 penetrates epithelial and endothelial cells and in most studies the electron micrograph images reveal a distribution of sizes and shapes for this bacterium (32). Despite the seemingly even distribution of bacterial sizes and shapes, the bacterial cells more closely associated with the epithelial tissue are spherical (32). Interestingly, under our experimental conditions, both IHE3034 and its ΔclbP mutant exhibited a uniformly spherical shape with none of the expected cylindrical shape, as observed by SEM. Some studies have suggested that alterations in the chemical composition of the peptidoglycan layers promote a rod to sphere change in morphology of some gram negative bacteria, an observation that could explain the presence of spherical *E. coli* (33). At present, based on the SEM images, the accumulation of colibactin in the wild type IHE3034 and the accumulation colibactin intermediates in the Δ*clbP,* in the periplasmic space, plays a role in altering the structure of the peptidoglycan layer causing this rod to sphere transition. However, further experiments may be needed to ascertain this effect by other characterization tools.

In conclusion, the successfully production and validation of an isogenic mutant of *E coli* IHE3034 without the gene encoding the *clbP* peptidase has been presented. The Δ*clbP* deletion had been carried out extensively in Nissle 1917 and in bacterial artificial chromosome (BAC) containing the *pks* genes (for further heterologous expression in a non-virulent strain) but never in the IHE3034 *E. coli* strain (11, 16).

Our results demonstrate that the envelope lesions observed in some *pks+* clinical isolates cannot be attributed to the presence of the *pks island* or to colibactin production. However, we observed an unexpected sphere shape in both IHE3034 and its isogenic mutant strain, which raises new questions about the biosynthetic pathway of colibactin and its interactions with the bacterial membrane before being exported out of the bacterial cell.

## Materials and Methods

### 1) Strains

*E. coli* clinical isolates, both *pks^+^* and *pks^−^,* were obtained as part of a of carbapenem-resistance from hospitals in Puerto Rico (34). *E. coli* IHE3034 *pks^+^* was kindly donated by Dr. Eric Oswald from the University of Toulouse, France. The *E. coli* DH10B *pks^−^* (non-genotoxic) was used in all the experiments as a control strain. All strains, including the Δ*clbP* mutant detailed later on, were “re-animated” from cryogenic storage in Luria broth (LB, Sigma-Aldrich) agar for 24 hours at 37 °C. Colonies were then selected and grown in liquid LB for 24 hours at 37°C. Followed by the corresponding experiments.

### 2) Deletion of the *ClbP* gene

Deletion of the *clbP* gene was achieved using Red/ET recombination following the Quick & Easy *E. coli* Gene Deletion Kit from Gene Bridges (Cat. No. K006). Kanamycin was used as the antibiotic selection marker. Primers with 5’-overhang of the first 50 bp of the *clbP* gene (homology arms) were used to amplify a lineal cassette (see Table 1). Briefly, overnight cultures of *E. coli* were prepared for electroporation. *E. coli* cells were centrifuged for 30 seconds at 11,000 rpm, the supernatant was removed, and the pellet was resuspended in sterile ultra-pure water (EMD Milllipore). This step was repeated two times. Then, cells were incubated with the pRed/ET plasmid encoding the recombinase for 1 minute. This plasmid contains the enzymes to perform the recombination between the *clbP*-homology arms of the lineal cassette and the *clbP* gene. This was followed by electroporation with 5 ms pulses of 1,350 V using the MicroPulserTM (BioRad). Cells were then resuspended in LB liquid (Sigma-Aldrich) medium for 70 minutes, at 30°C, and shaken at 100 rpm. The transformed *E. coli* cells were then plated in LB agar (Sigma-Aldrich) with ampicillin (Sigma-Aldrich) overnight. Lastly, pRed/ET+ *E. coli* colonies were selected and incubated in LB with ampicillin and 10% L-Arabinose for 60 minutes at 37°C to induce the expression of proteins necessary for recombination. The recovered cells were prepared for electroporation, incubated with the *clbP* homology arm-lineal cassette, and electroporated as described above. Cells were then incubated in LB for 3 hours at 37 °C and plated in LB agar with Kanamycin (Sigma-Aldrich) for 24 hours at 37°C. Colonies were selected and transfer to liquid LB, with 50 μL/mL of Kanamycin, for 24 hours at 37 °C and shaking at 250 rpm. The overnight culture was stored in glycerol at −80 °C.

**Table 1.**
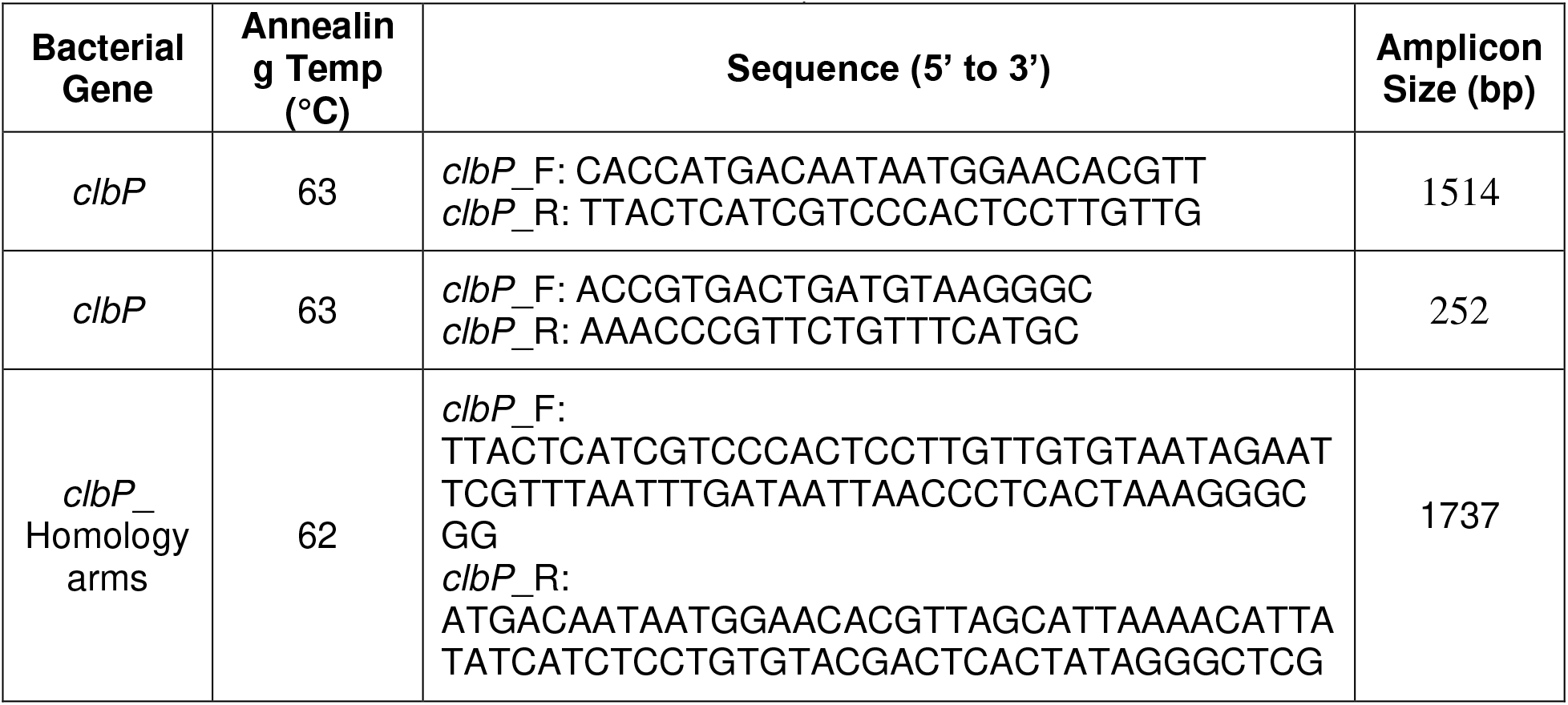
Summary of PCR conditions for amplification of the *clbP* flanked-lineal cassette and validation of the mutant strain *(ΔclbP)*

### 3) Confirmation of *ClbP* deletion by PCR

To validate that the *clbP* gene was successfully removed from the *E. coli* IHE3034 *pks^+^* strain, PCR was employed using genomic DNA from the Δ*clbP* strain. Specific primers that were used are summarized in Table 2. Briefly, using Phusion^®^ High DNA polymerase (NEB), an initial denaturation step of 1 minute at 98 °C was performed followed by 30 cycles of 10 seconds at 98 °C, 30 seconds at the corresponding annealing temperature (Table 2), and 2 minutes at 72 °C. All reactions were finalized with a final extension step of 10 minutes at 72 °C. *E. coli* DH10B *pks^−^* was used as a negative control, *E. coli* IHE3034 *pks^+^* as a positive control, and water as a PCR blank. All PCR products were visualized by running a 1% agarose gel electrophoresis stained with GelRed™(Biotium).

**Table 2.**
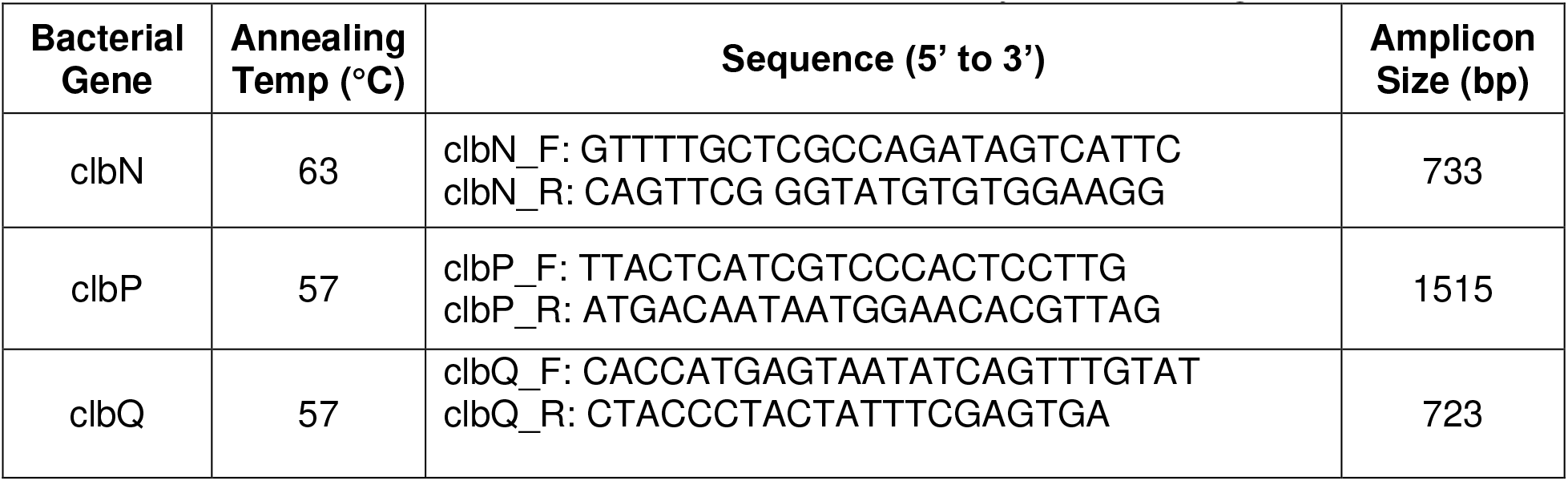
Primers used to confirm the deletion was solely of the *clbP* gene

### 4) Growth curves for IHE3034 and IHE3034 Δ*clbP*

Overnight culture of *E. coli* IHE3034 *pks^+^, E. coli* IHE3034 Δ*clbP,* and *E. coli* DH10B *pks^−^* were seeded in a 96-well plate in LB liquid medium (Sigma-Aldrich). The optical density (OD) was measured using the Synergy H1 hybrid reader (BioTek) at 600 nm for 24 hours at 37 °C under constant movement.

### 5) HeLa cells culture

HeLa cells (ATCC CCL-2) were cultured in Dulbecco’s Modified Eagle’s Medium (SIGMA Cat No. D5796) supplemented with 10% fetal bovine serum (SIGMA Cat. No. F2442) and Penicillin/Streptomycin (100 U:100 μg/mL) (CORNING Cat. No. 30-002CI). The cells were grown in a 24 wells culture plate with a total volume of 1 mL and with a sterile round coverslip in each well. They were then incubated at 37 °C, in 5% CO_2_, until 50% confluence was achieved to further infection.

### 6) Confirmation of *ClbP* deletion by HeLa infection

Approximately, 7.5 × 10^5^ bacterial cells were inoculated in 1 mL of DMEM with 10% FBS and without any antibiotic, this was then added to the HeLa cells and co-incubated for 4 hours. Then, the HeLa cells morphology was analyzed under a light microscope (Nikon ECLIPSE LV100N POL) using Giemsa staining, following the manufacturer’s instructions (SIGMA Cat. No. GS500). Briefly, cells attached to the round coverslip were fixed using methanol, followed by three consecutive washes with ddH2O. Then, cells were incubated at room temperature with 1 mL of 1:20 solution of Giemsa stain:water for 30 minutes. Coverslips were transferred upside down in an optical microscope slide with a minimal amount of Xylene based mounting media (Cat. # LC-A).

### 7) Scanning electron microscopy (SEM)

Samples for SEM analysis were prepared and processed as reported by Piroeva *et al.* Following is a brief description of each step (35).

#### Bacterial growth

Exponential growth was achieved overnight in LB medium at 37°C and 250 rpm. Bacterial samples were collected by centrifugation of the growth cultures at 3500 rpm for 5 min, followed by two pellet washes with ddH2O. Finally, 5–20 μL of sterile and filtered water were added and the pellet was carefully homogenized, thus ready for fixation for electron microscopy observation.

#### Sample fixation and dehydration

A coverslip (18 × 18 mm^2^), sterilized by UV irradiation for 10 minutes, was used as a platform for bacterial fixation. The coverslip was dipped in 0.8% agar solution and left horizontally, allowing the thin agar film to materialize for 30 minutes. Bacterial samples were carefully placed on the agar film and allowed to settle for 45 minutes. The samples were then dehydrated in an oven at 37 °C for 12 hours. Samples were processed by successive immersion in ethanol solutions from low to high concentrations (10, 25, 50, 75, 96 and 99 %). Samples were maintained in each ethanol solution for at least 30 minutes. Finally, samples were dried at 37 °C for about 1 hour.

#### Gold coating

Fixed and dehydrated samples are coated with a thin gold film (~10 nm) using a metal sputter system.

#### Scanning electron microscope (SEM) analysis

SEM analysis was performed with the high resolution field emission scanning electron microscope (JEOL JSM-7500F SEM), using 1 kV acceleration voltage for a gentle electron beam. The sample (coverslip) was mounted on a double coated conductive carbon tape that holds the sample firmly to the stage surface and a copper tape was placed as a ground strap from the sample surface to the SEM sample holder.

## Acknowledgements

This work was supported by National Institutes of Health (NIH) grant R25GM061151-15 (MBRS-RISE Program) fellowship to YML and grant R25GM061838 (MBRS-RISE Program) to RGM. Some of the shared instrumentation was purchased with NIH Grant U54 MD007600 (RCMI Program). The authors wish to thank Ms. Vilmarie Mercado and Dr. Yancy Ferrer for technical support and Dr. Carmelo Orengo, Dr. Rafael Ramos-Jiménez and Dr. Karylsa Torres for performing key critical feasibility experiments not included in this report.

